# Single-cell transcriptomes of fluorescent, ubiquitination-based cell cycle indicator cells

**DOI:** 10.1101/088500

**Authors:** Michael Böttcher, Tsukasa Kouno, Elo Madissoon, Efthymios Motakis, Imad Abugessaisa, Sachi Kato, Harukazu Suzuki, Yoshihide Hayashizaki, Takeya Kasukawa, Piero Carninci, Timo Lassmann, Jay W. Shin, Charles Plessy

**Author notes:** Corresponding authors: Michael Böttcher and Charles Plessy.

## Abstract

We used a transgenic HeLa cell line that reports cell cycle phases through fluorescent, ubiquitination-based cell cycle indicators (Fucci), to produce a reference dataset of more than 270 curated single cells. Microscopic images were taken from each cell followed by RNA-sequencing, so that single-cell expression data is associated to the fluorescence intensity of the Fucci probes in the same cell. We developed an open data management and quality control workflow that enables users to replicate the processing of the sequence and microscopic image data that we deposited in public repositories. The workflow outputs a table with metadata, that is the starting point for further studies on these data. Beyond its use for cell cycle studies, We also expect that our workflow can be adapted to other single-cell projects using a similar combination of sequencing data and fluorescence measurements.

## Background & Summary

Growing cells represent a heterogeneous population of cells alternating between different cell cycle phases. Several methods to investigate cell cycle stages are available, for instance fluorescence labeling of cell cycle specific marker proteins, staining with fluorescent DNA dyes or cell culture synchronization. However, these methods are often hard to combine with existing protocols for high-throughput single-cell gene expression analysis. The Fucci probe system is a set of fluorescent proteins fused with cell-cycle dependent degradation boxes[1]. In their progression through the cell cycle, Fucci cells alternate between red, green and no fluorescence. In short, only red fluorescence corresponds to G1, only green fluorescence to G2, a mix of both to some transitional stages from G1 to G2, and no fluorescence to some transitional stages from G2 to G1. We obtained fluorescence measurements of hundreds of individual Fucci cells (see methods) and generated unstranded paired-end RNA-sequencing libraries using the C1 microfluidics technology (Fluidigm). We intend to use the combined image and transcriptome data to enable the discovery of novel cell cycle marker genes that can be used together with known cell cycle related genes to infer cell cycle phases with single-cell expression data. In addition to this core data, we also collected metrics useful for quality control and curation: the cDNA yield after the first cDNA amplification, the sequencing yield, and the number of reads matching reference sequences such as spikes, ribosomal RNAs or known artefacts. The whole data processing workflow is implemented in literate programming based on R markdown files. We use three publicly accessible repositories to share all sequence and image data, as well as commented source code for data processing (Data Citation 1, 2, 3). A comprehensive overview of all experimental steps and data management is shown in Fig. 1.

**Fig. 1:**
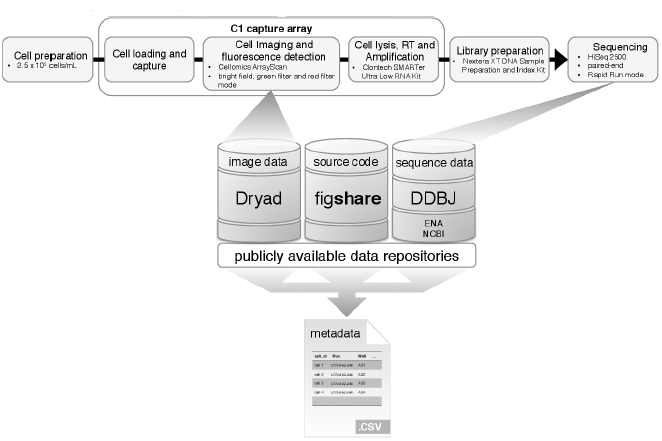
Experimental workflow and data management. We captured single cells via the C1 capture array and used an automated microscopy system to take images of all cell capture chambers. After imaging the cells were lysed, and cDNA synthesis as well as cDNA amplification were done in the Fluidigm’s commercial C1 Single-Cell Auto Prep System. Next, cDNA amplicons were transferred to a 96-well plate on which we utilized Nextera’s tagmentation technology to perform fragmentation and adaptor ligation simultaneously. After an index PCR step, single-cell samples were pooled and sequenced together on one lane of the Illumina HiSeq 2500 device. Image data in bitmap and C01 format were uploaded to the Dryad digital repository, however the DOI for the Dryad repository will only become publicly accessible upon publication in a peer reviewed journal. In the meantime the image data can be obtained elsewhere (Data Citation 2). Fastq sequence files were submitted to DDBJ. All scripts used to perform quality checks on raw data and to assemble a comprehensive metadata table for all single-cells are open source on figshare to ensure a high level of reproducibility.

## Methods

### Cell culture and capture

Cells were cultured in DMEM medium (Wako cat num. 044-29765) supplemented with 10 % FBS (Nichirei Biosciences, cat num. 171012, lot DCE7025) and penicillin/streptomycin (Wako cat num. 168-23191), and dissociated by trypsin treatment (Gibco). The cell size (between 14.1 and 15.2 *μ*m) and viability (92 to 98 %) of the cell concentration was estimated using the Countess Automated Cell Counter (Invitrogen) before loading 3,000 cells in a C1 single-cell Auto Prep arrays (Fluidigm, cat num. 100-6041) for mRNA-sequencing (10-17 *μ*m) following the manufacturer’s instructions. RNA spikes (ArrayControl RNA Spikes, Ambion, AM1780) were introduced in each chamber as described in the manufacturer’s user protocol. For the last three runs two more spikes (number 3 and 6) were added at a 10 and 100 times lower concentration respectively.

### Imaging

We used a Fucci variant, which contains mCherry, mVenus, and AmCyan, fused to the same regulatory domains of human Cdt1 and Geminin as used in the original Fucci [1], but in distict combination. Pictures of each cell captured in the C1 capture arrays were taken with a Cellomics ArrayScan VTI High Content Analysis Reader (Thermo Scientific). For each chamber, pictures were taken in bright field, with a green filter (excitation bandwidth: 480-495 nm, emission bandwidth: 510-545 nm), and with a red filter (excitation bandwidth: 565-580 nm, emission bandwidth: 610-670 nm) (Thermo Scientific). While these settings do not let us take advantage of the blue transgene, the recovered information is enough to order the cells on a pseudotime axis[2]. The full characterisation of the Fucci variant is beyond the scope of this work and will be presented in a separate manuscript that will be the primary reference for this new cell line. The quantification of fluorescence intensity was semi-automated using the free ‘Fiji’ software [3] with a customized macro (‘fluorescence’ folder [Data Citation 3]). The scanning and image acquisition process for one C1 capture array took about 15 min.

### Library preparation and sequencing

Single-cell RNA-sequencing libraries were prepared following the manufacturer’s manual (Fluidigm protocol PN 100-5950 A1). In brief, single-cell RNAs were reverse-transcribed with the SMARTer Ultra Low RNA Kit for the Fluidigm C1 System, (Clontech 634833/ TaKaRa Z4833N), the cDNAs were amplified inside the capture arrays with Advantage 2 PCR enzyme (Clontech, PN 639206), and transferred to a 96-well plate. They were then quantified by fluorimetry using the PicoGreen [4] intercalating dye and then fragmented by tagmentation with the Nextera XT DNA technology (Illumina FC-131-1096), followed by multiplexing by combination of 50 (S501-S508) and 30 (N701-N712) indexes (Illumina FC-131-1002) and sequencing (HiSeq 2500 Rapid Run mode, 150 nt, paired-end).

### Code availability

All scripts used for the processing of the sequence and image data are freely accessible on figshare (Data Citation 3). Custom scripts are available as R markdown files and are either written in R or as Linux shell scripts following the ‘literate programming’ and ‘reproducible research’ paradigms.

## Data Records

All data is deposited in three open repositories, for the source code and tables with raw data measurements, the sequencing output and the single-cell image files.

The RNA-sequencing paired-end fastq files can be accessed through the DNA DataBank of Japan (DDBJ) (Data Citation 1). A script for the batch download and renaming of fastq files according to unique cell identifiers instead of DDBJ accession numbers can be found in the ‘DDBJ’ folder on figshare (Data Citation 3).

Single-cell image files can be downloaded from or viewed directly using an inhouse developed database (Data Citation 2). Image files are available in bitmap and Cellomics system-specific C01 format. The C01 format files have been used for semi-automated quantification of fluorescence intensities. A detailed description on the intensity quantification and quality checks based on images can be found in the ‘fluorescence’ directory on figshare (Data Citation 3).

Apart from image and sequence data we also utilized raw data from fluorimetric assays to measure the cDNA concentration after the cDNA synthesis step in the C1 capture array, which can be found in the ‘cDNA_concentration’ folder on figshare (Data citation 3).

### Assembly of the metadata table

The metadata for each of the 480 sample observations, that is the final output of our workflow, is stored in a table called ‘combined.csv’ in the ‘combine_all’ folder on figshare (Data Citation 3). This table is to be used for quality assessment and downstream analysis of the dataset. It includes columns to identify cells with a unique ‘cell_id’, to indicate their coordinates in the 96-well plates used for tagmentation, to provide the normalized fluorescence intensities in the red and green filter channels and to indicate pass and fail results for quality checks such as the visual curation of cell images and the analysis of library read numbers. This table also identifies the standard positive and negative control samples, which were multiplexed in place of samples from chambers where no cell was captured. Lastly, this table includes columns with read counts matching reference sequences for spikes, rRNAs, Nextera primers and the HPV18 genome.

## Technical Validation

### Cell detection and image curation

All single-cell images were visually screened by two independent curators and cells were flagged with an error type of 0 if a normal cell was detected, 1 if no cell was present, 2 if debris was observed, 3 if the camera focus was off and 4 if more than one cell was found. When in doubt about it’s status, a cell was not flagged with type 0.

Initial inspection of the fluorescence values revealed some duplicates unlikely to happen by chance, which were the symptom of a software bug related to the case-insensitivity of the file system used to store the data. Cells affected by this data loss are flagged for removal in the metadata table, and the imaging procedure was corrected to avoid exhausting the alphabet letters when constructing the two-character file names used by the Cellomics system (the chambers in the C1 capture array are organized following a 48 × 2 geometry).

### Normalization of fluorescence intensites

For each fluorescence image we estimated the raw foreground spot intensity and the background intensity of a fixed cell-free area and summarized the quantified intensities by using the subtraction model [5]. Quality controls on the summarized channel signals highlighted their characteristic multi-modal distribution and the existence of run-specific, non-biological variation that had to be corrected prior to further analysis. Among others, we assume that small differences in exposure time settings for the first two runs (1772-062-248 and 1772-062-249) are a source of the observed variation. Furthermore, an auto-focus option was used for the first three runs, whereas fixed camera settings have been applied to the last two runs (1772-067-038, 1772-067-039).

We fitted the run factors on the log-transformed background-adjusted signals of each channel and applied a linear one-way ANOVA model to remove the effects from the data. This strategy had two drawbacks. First, the range and the modes of the background-adjusted signal distributions differed substantially across runs (potentially as a result of the image capture settings). Thus, applying one-way ANOVA simply shifted the data distributions to equalize their means, but it did not adjust for run effects. Second, the model does not satisfy the N(0, *σ*^2^) assumption for the residuals. The respective P-value of the Kolmogorov-Smirnov test for normality was less than 1%.

To detect and remove the unwanted effects, we fitted the log-transformed signals of each channel to a latent class, Bayesian mixture of regression model [6, 7] that estimated the optimal number of mixtures (modes of the signal distribution) for each run, and removed the run effect by a two-way factorial ANOVA model. The factors of that model were the runs, the estimated mixtures (a binary variable separating the sub-populations around each mode) and their interaction. This model produced comparable signal densities across runs and generated approximately N(0, *σ*^2^) distributed residuals. The run corrected signals were consequently adjusted by the normexp model [8], that deconvolved the signal into its exponentially distributed component (representing the estimated spot intensity) and its normally distributed component (accounting for random noise). By design, the estimated signals of the normexp model are bound to be positive (it sets all negative background-adjusted, run-corrected signals into small positive values changed by a constant k that minimizes the data variability [8]). The possibility of having negative background-adjusted, run-corrected signals in both channels was excluded by our quality filters. However, negative values in a single channel are possible, biologically meaningful, and eventually associated to a certain cell cycle phase. All scripts and plots for our fluorescence data correction method can be found in the ‘Intensity_correction’ directory on figshare (Data Citation 3).

### cDNA yields

Within the C1 system cells are lysed, polyadenylated RNAs are reverse-transcribed and cDNAs amplificated by PCR. These PCR products are then transferred to a 96-well plate (one well per cell) and their concentration is quantified by fluorimetry. We observed a strong variation of the average yield among the five C1 capture array runs (Fig. 2a). Nevertheless, except for the 3rd run 1772-064-103, where the concentration never exceeded 0.7 ng/*μ*L, a bimodal distribution of cDNA yields can be seen (‘cDNA_concentration’ directory [Data citation 3]). Since every chamber contained RNA spikes no yield was exactly zero, but the lowest yields correspond to chambers of the C1 capture array where no cell was captured (Fig. 2b). This suggests that in situations where it is not possible or desirable to take images of each chamber, the cDNA yields may be used to flag chambers of the C1 capture array that had no cell.

**Fig. 2:**
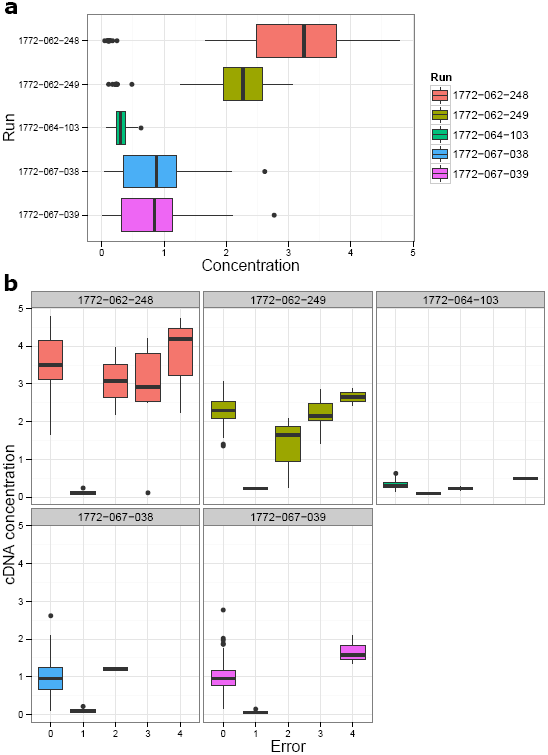
Distribution of cDNA concentrations. Each Run represents 96 sample concentration measurements [ng/*μ*L], including 2 controls per Run. 480 samples in total for all five Runs. Individual Runs are technical replicates, whereas most samples within a Run are biological replicates. Distributions of sample measurements are displayed as boxplots colored by Run (**a**, **b**). The top panel (**a**) Shows the cDNA concentration per Run and the bottom panel (**b**) shows for each Run how sample concentrations are distributed among four different error types. These error types are: 0 = cell present; 1 = cell absent; 2 = debris present; 3 = wrong focus; 4 = more than 1 cell. ((**a**) ‘cDNA_concentration’ and (**b**) ‘combine_all’ directory on figshare [Data citation 3])

### Sequencing

After tagmentation and multiplexing with the Nextera kits, single-cell libraries were pooled and sequenced in Rapid Run mode on HiSeq 2500 sequencers. Individual runs were sequenced on separate flow cell lanes, yielding between 172,535,190 and 110,618,416 read pairs. The number of pairs per cell was therefore higher than 1 million on average. Given that the final steps of the library preparation before multiplexing were done in 96-well plates, we inspected the sequencing yields for possible bias correlating with rows or columns of the plates (Fig. 3a, b) (‘HiSeq’ directory [Data Citation 3]). Although no plate position bias was found, we detected a probable pipetting error, where libraries from column 08 of the plate in run 1772-064-103 had almost twice as many reads as the previous columns of the same plate. Additionally, libraries from its neighbor column 09 did not yield a considerable read number (Fig. 3b). The single-cell libraries corresponding to these two columns of run 1772-064-103 were consequently flagged for removal.

**Fig. 3:**
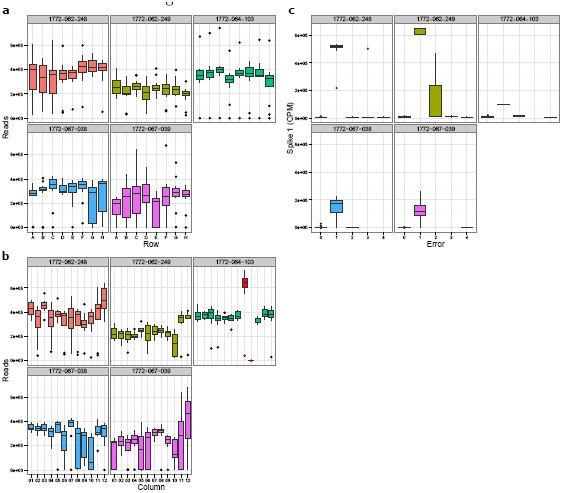
Read count distributions. The distribution of sequencing reads of cells grouped by their position in rows (**a**) and columns (**b**) on the 96-well plate used for sequencing library preparations. The data points of column 08 and 09 in the third Run (**b**) are highlighted in red to mark them as outliers created by a pipetting error. The right sight panel (**c**) shows Spike 1 counts per million of total reads (CPM) grouped by error type. ((**a**) and (**b**) ‘combine_all’ and (**c**) ‘HiSeq’ directory on figshare [Data citation 3])

### RNA spikes and control sequences

Following Fluidigm’s protocol, a serial dilution (10*x*) of ArrayControl RNA spikes (Ambion) was present in the cell loading buffer, and therefore at an equal concentration in each chamber. Using the TagDust2 software [9], we counted the number of reads matching the spike sequences (‘control-sequences’ folder [Data Citation 3]). In absence of cells in the capture chamber, the proportion of spikes in the libraries was strongly elevated (Fig. 3), which suggests another possible way to detect the absence of cells when no microscopy images are available. Using the same approach, we also counted the reads matching rRNA, Nextera primers, and the HPV18 genome, of which a fragment is integrated in the HeLa cells.

## Usage Notes

The dataset can be used to identify known and novel transcript markers of the cell cycle. One aim of the downstream analysis of this dataset is to create a model for cell cycle status inference on the single-cell level based on expression data. Previously, a new method to identify transcriptional dynamics of oscillating genes in single-cell RNA-seq data was tested using H1-Fucci cells among others [10]. Other groups have used single-cell transcriptome data in mouse embryonic stem cells to estimate the expression variance contributed by the cell cycle[11], and in different human cell lines to estimate the amount that cell cycle contributes to overall variability of gene expression in individual cells [12]. Nevertheless, no reliable cell cycle prediction model for single-cells has been established yet. Moreover, there are other possible uses, such as the study of cell cycle specific splice forms (taking advantage of the 150 nt read length) or the inference of regulatory networks underlying cell cycle phase transition events, or simply for benchmarking of other C1 RNA-sequencing datasets as well as analysis workflows. There are a few limitations with this dataset. We expect batch normalization to be necessary because the data was produced over 4 months by multiple operators on different sequencing instruments. Three out of five runs have much reduced spike concentrations, therefore the spike data in this dataset cannot be used for data normalization (‘control-sequences’ folder [Data Citation 3]). Also, our experimental design is not suited for the study of the G0 phase. In future works, this issue might be addressed by using primary cells from Fucci mouse [13] or Fucci zebrafish [14].

The analysis workflow was written and documented in order to be reusable in other studies. In particular, the default C1 RNA-sequencing workflow includes a live/dead staining that also produces red and green fluorescence values, which fits into our pipeline.

An integrated single-cell database platform is developed within RIKEN CLST and can be used to quickly screen image files and metadata of the HeLa Fucci dataset (Data Citation 2).

## Acknowledgements

This work was supported by a Research Grant from the Japanese Ministry of Education, Culture, Sports, Science and Technology (MEXT) for RIKEN Omics Science Center from MEXT to YH and to the RIKEN Center for Life Science Technologies, a Grant-in-Aid for Young Scientists A (number 25710018), and by the RIKEN International Program Associate program. The authors wish to acknowledge Asako Sakaue-Sawano and Atsushi Miyawaki for providing the Fucci cell line prior publication, Mami Kishima for cell culture, RIKEN GeNAS for the sequencing of the libraries using the HiSeq 2500, and Fumi Hori for data deposition to DDBJ.

Author contributions: Designed the single-cell transcriptome analysis of Fucci cells: PC, TL, JS, CP. Planned transcriptome analysis of Fucci cells: HS, YH. Developed fluorescence acquisition workflow: TK, JS. Produced RNA-seq and fluorescence data: TK, SK, JS. Developed fluorescence quantification macro: EloM. Quantified fluorescence data: MB, EloM. Developed and applied the normalization strategy for the fluorescence data: EM. Built database for image and metadata storage: IA, TaK. Integrated the data: MB, TL, JS, CP. Implemented the R markdown workflow: MB, CP. Wrote the manuscript: MB, CP.

